# ExomeSlicer: a resource for the development and validation of exome-based clinical panels

**DOI:** 10.1101/248906

**Authors:** Rojeen Niazi, Michael A. Gonzalez, Jorune Balciuniene, Perry Evans, Mahdi Sarmady, Ahmad N. Abou Tayoun

## Abstract

Exome-based panels (exome slices) are becoming the preferred diagnostic strategy in clinical laboratories, especially for genetically heterogeneous disorders. The advantages of this approach include enabling frequent updates to gene content without the need for re-designing, reflexing to exome analysis bioinformatically without requiring additional sequencing, and streamlining laboratory operation by using established exome kits and protocols. Despite their increasing use, there are currently no guidelines or appropriate resources to support their clinical implementation. Here, we highlight principles and important considerations for the clinical development and validation of exome-based panels, guided by clinical data from a diagnostic epilepsy panel using this approach. We also present a novel, publically accessible web-based resource, ExomeSlicer, and demonstrate its clinical utility in predicting gene-specific and exome-wide technically challenging regions that are not amenable to Next Generation Sequencing (NGS), and that might significantly lead to increased post hoc Sanger fill in burden. Using this tool, we also characterize > 2000 low complexity, GC-rich and/or high homology, regions across the exome that can be a source of false positive or false negative variant calls thus potentially leading to misdiagnoses in tested patients.

## Introduction

Next Generation Sequencing (NGS) has proven to be a powerful tool for the identification of genetic variants in Mendelian disorders, and as such played major roles in disease gene discovery in research setting and also clinically for establishing genetic diagnoses [1–3]. While sequencing of the entire genome (whole genome sequencing [WGS]) represents the most comprehensive option, sequencing of coding regions only, so-called whole exome sequencing (WES), is the cheaper and more practical alternative [4].

Using WES or targeted NGS panels as a clinical platform has proven to achieve high, up to 50%, diagnostic yield for a range of clinical indications [5–10]. Several factors affect the decision regarding a suitable sequencing approach. While WES is reserved for complex clinical presentations, or for highly suspected genetic etiologies where exhaustive targeted testing did not reveal answers, NGS gene panels are mostly used for patients with rather simple clinical presentations or specific syndromes associated with a number of known disease genes [5].

Despite the advantages of WES, there are some drawbacks associated with this approach including its high cost, long turnaround time besides its analytical and interpretative challenges. WES uncovers numerous variants, including many of uncertain clinical significance, besides others that are unrelated to patient’s indication raising ethical concerns for the patient and their families [11, 12]. On the other hand, targeted NGS panels yield a very limited number of variants in the captured genes only, and with a relatively fast turnaround time. However, given the *priori* overlap between a patient’s phenotype and diseases caused by genes on the panel, a complete test, wherein regions with low depth of coverage are filled in using an orthogonal – most often Sanger sequencing – approach, is highly warranted [7, 13]. A major limitation of targeted NGS approach, however, is the static nature associated with its fixed number of captured genes. Thus, newly discovered genes cannot be easily added to the test and would require significant design and validation efforts by the laboratory.

The exome-based targeted panel approach (also called “Exome Slice”) has therefore recently become more attractive to laboratories as it allows dynamic gene content update with minimal design and validation, and at the same time reduces the interpretation burden associated with WES. This is most suitable for diseases that are known for high genetic and phenotypic heterogeneity such as epilepsy and hearing loss where at least a hundred gene might be involved [14, 15]. In this approach, the exome is sequenced but only indication-relevant genes are analyzed and interpreted using *in silico* bioinformatics tools. For laboratories offering exome sequencing and targeted gene panels, this approach can significantly homogenize the upfront analysis leading to a uniform wet-bench exome capture workflow. In addition, it allows for reflexing to exome analysis bioinformatically without requiring additional sequencing.

However, exome-slices are subject to the same downsides of exome captures including potential lack of adequate coverage for all the relevant genes. Additionally, more bioinformatics analysis is needed during test development and validation to identify gaps in coverage and/or regions of high homology and to design ancillary assays if warranted. Currently, there are no uniform standards or guidelines for the development and validation of exome slices in a clinical setting.

Here, we highlight important considerations for the clinical development and use of exome-based panels. We also develop a user-friendly, web-based tool, ExomeSlicer, for the identification of gene-specific and exome-wide technically challenging regions that cannot be reliably sequenced and that can be a source of false positive and/or false negative variant calls. We show that this tool supports exome-based and targeted panel development through identification of potential ancillary testing, characterization of test limitation, and streamlining post hoc Sanger sequencing.

## Materials and Methods

### Exome Capture and Sequencing

Exome-wide capture of the coding regions was performed using the SureSelectXT Clinical Research Exome (CRE) V5 kit (Agilent, CA, USA), followed by cluster generation using the TruSeq Rapid Cluster Kits (Illumina, CA, USA), and then sequencing on the Illumina HiSeq 2500 platforms with 2x100bp paired-end reads and average sequencing depth of 100X.

### Targeted Next Generation Sequencing

A custom-made capture kit was designed using RNA oligonucleotides targeting all RefSeq coding exons (± 25bp) of 338 disease genes (**Supplementary Table 1**) at 3X tiling density (Agilent SureDesign, Santa Clara, CA). Captured samples were prepared for sequencing using Agilent’s SureSelect QXT Library Preparation kit and sequenced using Illumina MiSeq or HiSeq 2500.

### Sanger Sequencing

M13-tagged primers were custom designed for the specific low coverage regions (see below) using an in house primer design software. Amplified DNA products were first cleaned to remove excess primers and dNTPs, and then sequenced using BigDye Terminator Cycle Sequencing Ready Reaction Kit (Applied Biosystems, CA, USA). Fuorescence-based cycle sequencing was performed using ABI 3730 automated DNA sequencer (Applied Biosystems, CA, USA). Sequences were viewed using Mutation Surveyor (SoftGenetics, PA, USA).

### Bioinformatics

Bioinformatics analysis of exome data was performed using the in-house “cwes-2.x” pipeline. Raw data (FASTQ) files generated by HiSeq were processed through this several components of this pipeline including hg19 reference genome (human_g1k_v37), Novoalign v3.04.06 (for alignment), SAMtools v1.1 (to convert SAM files to BAM files), Picard v1.123 (to mark duplicate reads and calculate coverage statistics), GATK v3.6 (for variant, SNV and INDEL, calling), and Snpeff v4.2 (for variant annotation). The pipeline generates several files (including VCF, annotations, quality report etc) however, for ExomeSlicer, we used BAM file after mark duplicate step. The same above components have been utilized for NGS panels.

### ExomeSlicer

Mapped reads BAM files from 1932 exome samples were used to calculate coverage and mapping quality statistics for ExomeSlicer. To calculate exon level coverage and mapping quality statistics for each sample’s bam file (after marking duplicate reads), Refseq coding exon hg19 coordinates, transcript and gene annotations were downloaded from UCSC (March 26, 2017). Using CRE capture bed file, each exon was annotated with number of baits covering the exon and the number of bases covered by baits. We developed a custom algorithm using PySam v0.12.0 to calculate statistics for mapping quality and coverage per exon (**Figure 1**). Coverage calculation was done by counting only “usable coverage” defined as: For a given base pair, only reads with Base Quality (BQ) ≥ 20 and Mapping Quality (MQ) ≥ 20. The MQ and BQ thresholds for counting coverage were used to ensure only reads contributing to variant calling were considered as most variant callers as recommended by Sanghvi *et al* [16]. For each sample, minimum, maximum, and average coverage all bases within an exon were calculated. Similarlyminimum, maximum, and average mapping quality of all reads covering an exon were calculated. To aggregate data across 1932 samples, average of each data point for each exon was calculated across all samples (e.g. average of all minimum coverages for a given exon across 1932 samples) as depicted in **Figure 2**.

**Figure 1.**
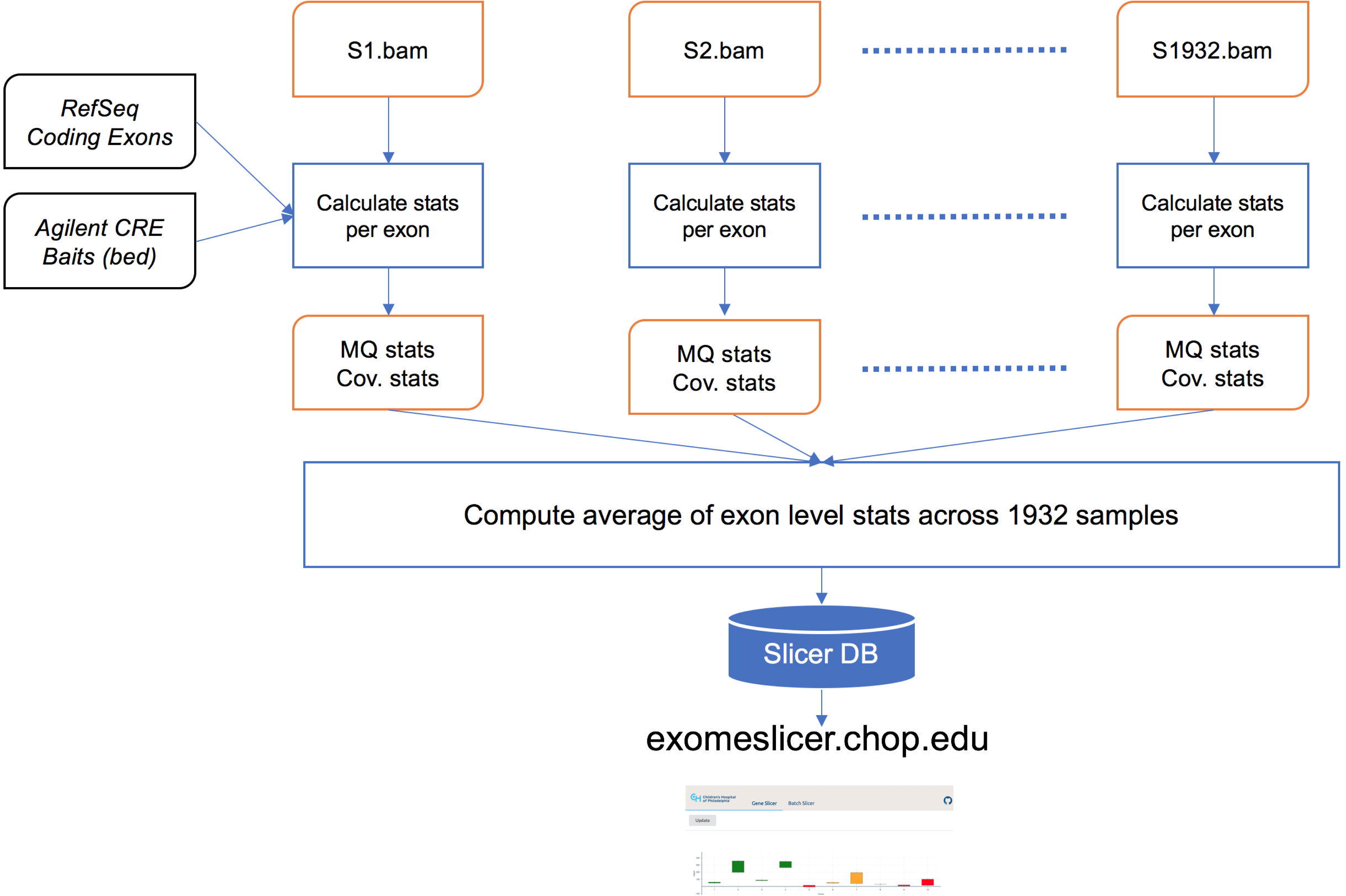
Exome Slicer data generation and visualization process. Exon level statistics were calculated per sample. Sample level data were then used to calculate aggregate statistics across 1932 clinical whole exome samples used to generate the ExomeSlicer Database. A web application was designed (exomeslicer.chop.edu) to visualize data from the database in an interactive way.

**Figure 2.**
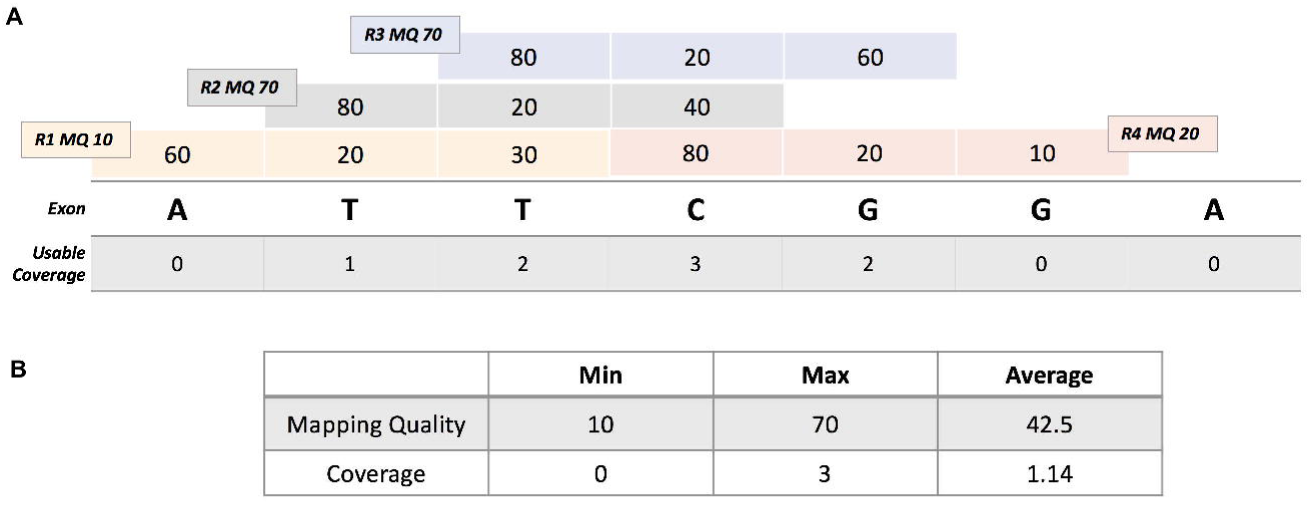
An example of ExomeSlicer exon level calculation. **A**, Seven base exon covered by four reads R1, R2, R3, R4. R1 is not counted in coverage of any of the bases because it does not satisfy MQ ≥ 20 condition of “usable coverage” as defined in methods. Similarly, last base **G** has zero coverage as the only read covering the base has BQ < 20. **B**, Statistics table for the exon. Average mapping quality is calculated using all reads (R1 through R4). Average coverage is calculated using coverage of each exon base.

### Statistical analysis

Proportions of good and poor quality regions with different baited bases were compared using a two-sample t-test. For additional region comparisons, overlapping regions were merged using bedtools merge. Merged region GC content was taken as the minimum of combined regions. BLAT hits for merged regions were found by running BLAT with default parameters against hg19. Merged regions were annotated with disease mutations from the Human Gene Mutation Database version 2016 and pathogenic, likely pathogenic, and variants of uncertain clinical significance from ClinVar (version February 2016) using bedtools intersect. Merged region genes were located in the Online Mendelian Inheritance in Man (OMIM) database (version 2013) using the OMIM genemap file. The GC content and number of BLAT hits for regions were compared using a two-sided Wilcoxon test.

## Results

### Development and Validation of Clinical Exome-based panels

#### Gene Content

The first step in the development of a sequencing panel is defining the genes wherein sequence variants are an established cause of the disease of interest. Although not specific to exome panels, this step is most critical in this approach given the larger number of genes that are likely to be included relative to the targeted capture approach. Therefore, consistent and robust selection criteria are needed to avoid inclusion of genes with limited or no clinical validity that could lead to unnecessary extra downstream analysis and interpretation, in addition to increased patient anxiety associated with added reporting of variants of uncertain clinical significance [14]. The Clinical Genome (ClinGen) Resource recently developed an evidence-based framework for evaluating the clinical validity of gene-disease associations [17]; using this framework is highly recommended for optimal gene content decisions in clinical exome-based panels.

Another important consideration is that the phenotype(s) caused by the selected clinically valid genes on a panel should be significantly overlapping or is/are at least the most prominent feature(s) associated with those disease genes. This especially critical for highly heterogeneous genetic disorders, which are most amenable to the exome-based approach, with several contributing genes, each can be associated with a varied phenotypic spectrum. For example, sequence variants in over 400 genes cause seizures or epilepsy besides several other features [15]. However, to design a sequencing panel for the specific diagnosis of epilepsy, then clinically valid genes known to cause pure epilepsies (non-syndromic or apparent non-syndromic) should only be included. Around 100 genes would now satisfy these inclusion criteria (**Supplementary Figure 1**).

#### Exome Capture

The most important consideration associated with using an exome capture is whether the relevant disease genes have adequate sequencing coverage or not. With the large number of targets in an exome, the overall coverage is expected to be low, as compared to smaller capture approaches, thus potentially leading to sequencing gaps. However, several commercial exome kits have now boosted the number of baits targeting coding exons, especially in disease genes, leading to adequate coverage across [18]. For example, in collaboration with Agilent and researchers from Emory University, we have recently developed an exome capture kit, the Agilent SureSelectXT Clinical Research Exome (CRE) V5 Plus kit, through which 96% of the coding regions (and even higher for disease genes) are covered with at least 10x read depth when 75M reads are sequenced [18].

To illustrate the efficacy of the CRE V5 plus kit for exome-based panels, we designed a different custom capture kit targeting 338 genes well known to cause a range of disorders including cholestasis, pancreatitis, lung disease, connective tissue disease, Cornelia de Lange, primary ciliary dyskinesia, and Noonan among many others (**Supplementary Table 1**). We then assessed coverage across the 338 genes when sequenced after the targeted custom capture or the CRE kit. As shown in **Supplementary Figure 2**, coverage across all genes is almost identical between the exome and the targeted custom capture kits (coding regions with at least 10x coverage was 98.6% and 98.9%, respectively) despite sequencing to an overall coverage of 100x and 400x, respectively. These data clearly demonstrate the utility of using current exome capture kits, such as the Agilent CRE, as a platform for the development of clinical exome-based panels.

#### Analytical Performance

A reference sample is needed as a benchmark to evaluate variant detection performance of NGS-based assays [13]. Genomic DNA of the HapMap sample NA12878 has been extensively sequenced and studied using multiple sequencing platforms in laboratories across the world. Moreover, the National Institute of Standards and Technology-Genome in a Bottle (NIST-GIAB) consortium has curated highly confident sequence variants across the whole genome of NA12878 and catalogued a benchmark reference variant dataset for this sample, including 28,940 single nucleotide variants (SNVs) and 996 small insertions or deletions (Indels) [19]. Using the CRE kit and sequencing the NA12878 sample at 100x read depth, the analytical sensitivity, specificity and positive predictive value for detecting the high confidence variants were 99.6%, 99.9%, 99.9%, respectively. This analytical performance further highlights the efficacy of using an exome-based approach for disease panels.

To assess the performance characteristics of the exome-based approach in any disease genes of interest, the high confidence NA12878 NIST-GIAB variants calls in this gene subset can be extracted and analyzed. For example, within the 100 epilepsy genes in **Supplementary Figure 1**, a total of 110 NIST-GIAB variants (including 108 SNVs and 2 indels) are found in the NA12878 sample. The calculated analytical sensitivity, analytical specificity, and positive predictive value in the epilepsy region of interest (~432kb) using the CRE kit with 100x read depth are 99.09%, 99.99%, and 99.09%, respectively. Such analysis complements the coverage information and ensures highly acceptable variant detection in the genes of interest.

#### Filtration

Upon exome sequencing, a validated filtration strategy is needed to retain rare variants only in the disease genes of interest. Removal of putative benign variants is critical to reduce the interpretation burden. This requires carefully setting a filtering allele frequency cutoff based on known disease attributes such as prevalence, penetrance, genetic and allelic heterogeneity, and mode of inheritance. Using variant frequency information in large representative individuals from the general population, such as the Exome Aggregation Consortium or ExAC [20], the diseases-specific cutoff can now be used to remove “common” benign variants.

Several conservative assumptions has to be made when calculating a disease-specific filtering cutoff due to lack of information regarding the above disease attributes. For example, the *SCN1A* gene, the most common cause of autosomal dominant (AD) epilepsy, can be used to establish a filtering cutoff for all AD epilepsy genes (**Supplementary Figure 1**). This gene has an estimated prevalence of 1/20,000 with a penetrance in the 70-90% range [21, 22]. Assuming that a single pathogenic variant causes all *SCN1A-related* seizures and that disease prevalence and penetrance are 1/10,000 and 50%, respectively, a conservative filtering cutoff of 1 X 10^−4^ for AD epilepsy is calculated using the Hardy-Weinberg equation or the recently published calculator (https://cardiodb.org/allelefrequencyapp/) [23]. Before clinical application, the filtration strategy has to be validated using samples with known pathogenic variants.

### Sanger Sequencing Burden

Despite the high overall coverage that can be achieved using a targeted or an exome-based capture approach (**Supplementary Figure 2**), certain regions might still fall below the required coverage for reliable variant detection. Such sequencing gaps are more likely to occur in regions with sequence complexities, such as GC-rich, repeat, or high homology sequences, which might affect target capture and/or amplification. It is expected that, for disease panels, a laboratory would make every effort to complete sequencing of all disease genes [7]. In fact, most clinical labs use Sanger sequencing to fill in regions falling below a certain coverage cutoff (usually below 15 or 20x). Given its post hoc nature, this process is cumbersome and can significantly delay the delivery of test results.

To illustrate this burden, around 1000 Sanger sequencing reactions were performed to fill in coverage gaps (defined as any region with at least 10bp with less than 15x) in 100 epilepsy cases originally tested using an exome-based approach shown in **Supplementary Figure 1**. Interestingly however, most (>900) of those Sanger reactions were recurrent (**Figure 3**) with 8 regions filled in ~700 times (**Table 1**). As expected, all 8 regions were either GC-rich or had repeat or high homology sequences (**Table 1**).

**Figure 3.**
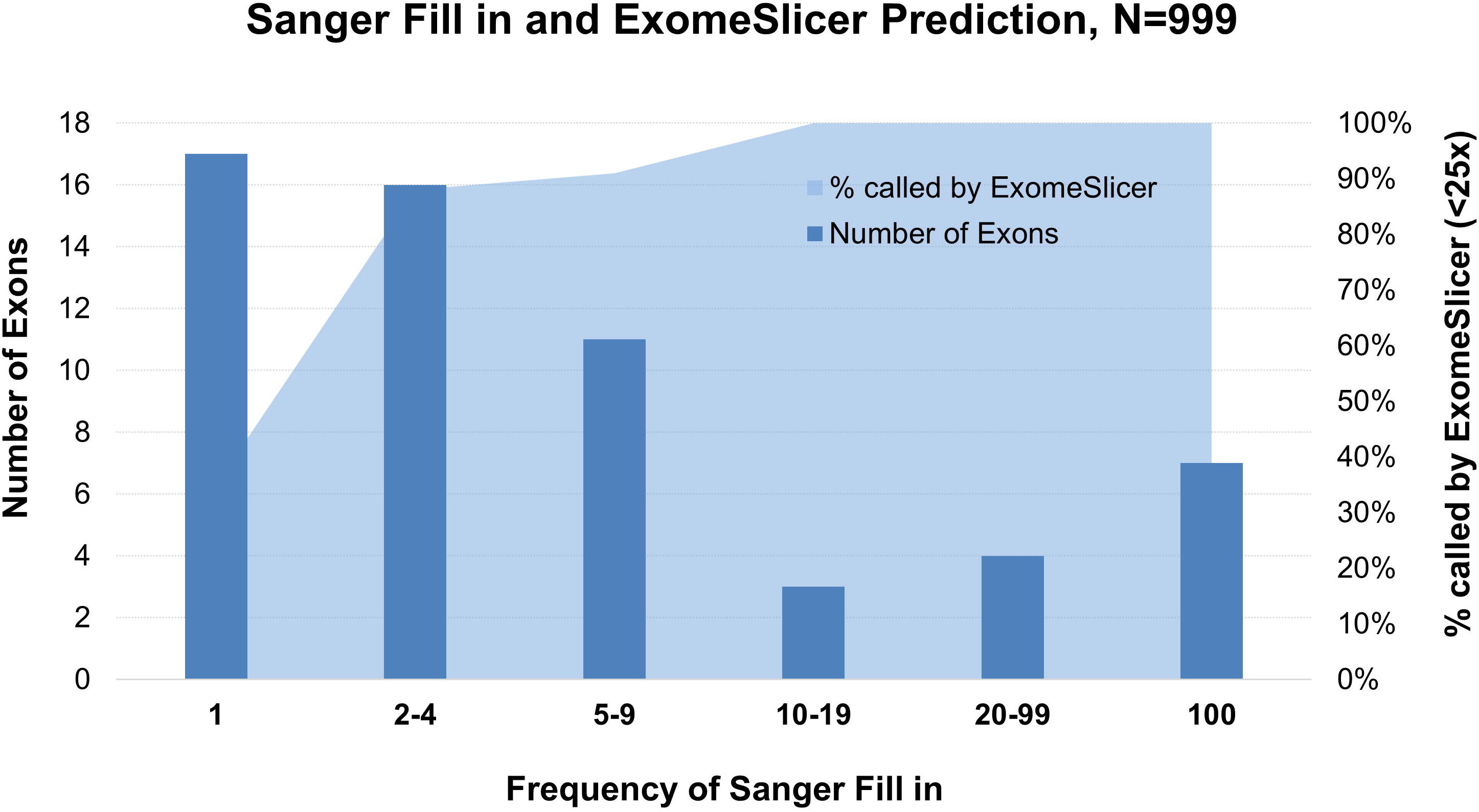
Post hoc Sanger sequencing fill in performed in 100 epilepsy cases tested using an exome-based approach. Most reactions were recurrent (frequency of fill in > 2) and were predictable by ExomeSlicer when the minimum average DP is at < 25x. Singleton exons, filled in only once (n=17), were sporadic and thus were not as predictable by ExomeSlicer.

**Table 1.** Technically challenging regions in epilepsy genes

| Gene | Transcript | CDS Exon | % Baited | Avg MQ | Min DP | ExomeSlicer Warning | Assessment |
| --- | --- | --- | --- | --- | --- | --- | --- |
| **ARX** | NM_139058 | 2 | 98% | 69.69 | 4.92 | Coverage | Poly-Alanine repeat region |
| **CERS1** | NM_021267 | 1 | 81% | 69.93 | 2.52 | Coverage | GC-rich |
| **EPM2A** | NM_005670 | 1 | 100% | 69.8 | 6.8 | Coverage | GC-rich |
| **IQSEC2** | NM_001111125 | 15 | 89% | 68.98 | 3.37 | Coverage | Proline and Histidine repeats |
| **SYN1** | NM_133499 | 12 | 94% | 69.61 | 8.14 | Coverage | GC-rich |
| **WDR45** | NM_001029896 | 10 | 37% | 69.82 | 5.11 | Coverage | End of exon with Poly Asparates |
| **SLC6A8** | NM_001142805 | 1 | 96% | 69.85 | 8.55 | Coverage | GC-rich |
| **SLC6A8** | NM_001142805 | 3 | 44% | 61 | 8.24 | Homology | Significant homology to other genomic regions |

CDS: Coding Sequence; Avg MQ: Average Mapping Quality; Min DP: Minimum Depth

Given the Sanger burden introduced by such regions, we tested whether we can identify them using exome sequencing data from ~2000 clinical samples (see methods and **Figures 1** and **2**). Highlighting such regions in the exome panel (or other targeted) approach can minimize the downstream post hoc analysis by designing separate upfront specific ancillary assays or by acknowledging them as part of the test limitation. We hypothesized that regions with sequencing complexities as in the above can be identified using NGS quality metrics, namely exon-level mapping quality (MQ) and depth of coverage (DP). In fact, all the regions in **Table 1** consistently had average minimum coverage below <10X and for one exon (*SLC6A8* exon 3) also had an average MQ score less than the optimal “70” score. Furthermore, most recurrent Sanger filled in regions are identified if a 25x, instead of 10x, minimum coverage is used (**Figure 3**).

Finally, it is worth noting that out of ~1000 Sanger fill in of low coverage (<15x) regions (**Figure 3**), no variants were detected that would have otherwise been missed by NGS. We therefore strongly recommend lowering the fill in cutoff (to <10x for example) and limiting to regions in genes with strong clinical validity, and wherein disease causing variants have been previously reported.

### Resource for Exome-based and Targeted Panel development

#### ExomeSlicer

Based on our analysis above and the potential utility for genes other than epilepsy, we developed a tool, ExomeSlicer (http://exomeslicer.chop.edu/), to support exome-based panel development through identification of technically challenging regions in any gene or gene lists across the exome. This tool uses empirical NGS quality metrics generated from 1932 samples that underwent clinical exome sequencing (**Figures 1** and **2**), and enables users to select appropriate key exon-level quality metric cutoffs and relevant transcript(s) to view or download technically challenging regions in their disease genes of interest. As shown in **Figure 1**, ExomeSlicer calculates average minimum, average maximum, and mean average MQ and DP in each baited exon across the exome. These values can be viewed using the “Gene Slicer” functionality, which provides an exon-by-exon MQ or DP plots for any queried gene/transcript, in addition to a downloadable table containing annotated exons and their corresponding quality metric values. Furthermore, users can provide a list of genes in the “Batch Slicer” functionality to retrieve a downloadable file with poor regions based on quality metrics that they can interactively provide.

#### Utility of ExomeSlicer

We chose mean average MQ and average minimum DP as key metrics for low coverage and/or high homology regions. Since any given exon can randomly have reads with a range of very low or very high MQ scores, we picked the mean average MQ value as a more reliable metric for unique mapping/homology issues. The average minimum DP was chosen to highlight low coverage regions, due to GC-rich or repeat sequences, not only in the whole exon but also “within” an exon, which can be missed if the mean average coverage was used instead. Using an average MQ and minimum DP cutoffs of 20 each [16], Gene Slicer detected known technically challenging sequences, including an intragenic tandem triplication in *NEB* [24], a pseudogene in *STRC* [25], a GC-rich exon in *SLC6A8* [26], and a Poly-Alanine repeat within exon 2 of *ARX* [27] (**Figure 4**). Furthermore, most epilepsy low coverage regions requiring Sanger fill in were predicted by ExomeSlicer (**Figure 3**).

**Figure 4.**
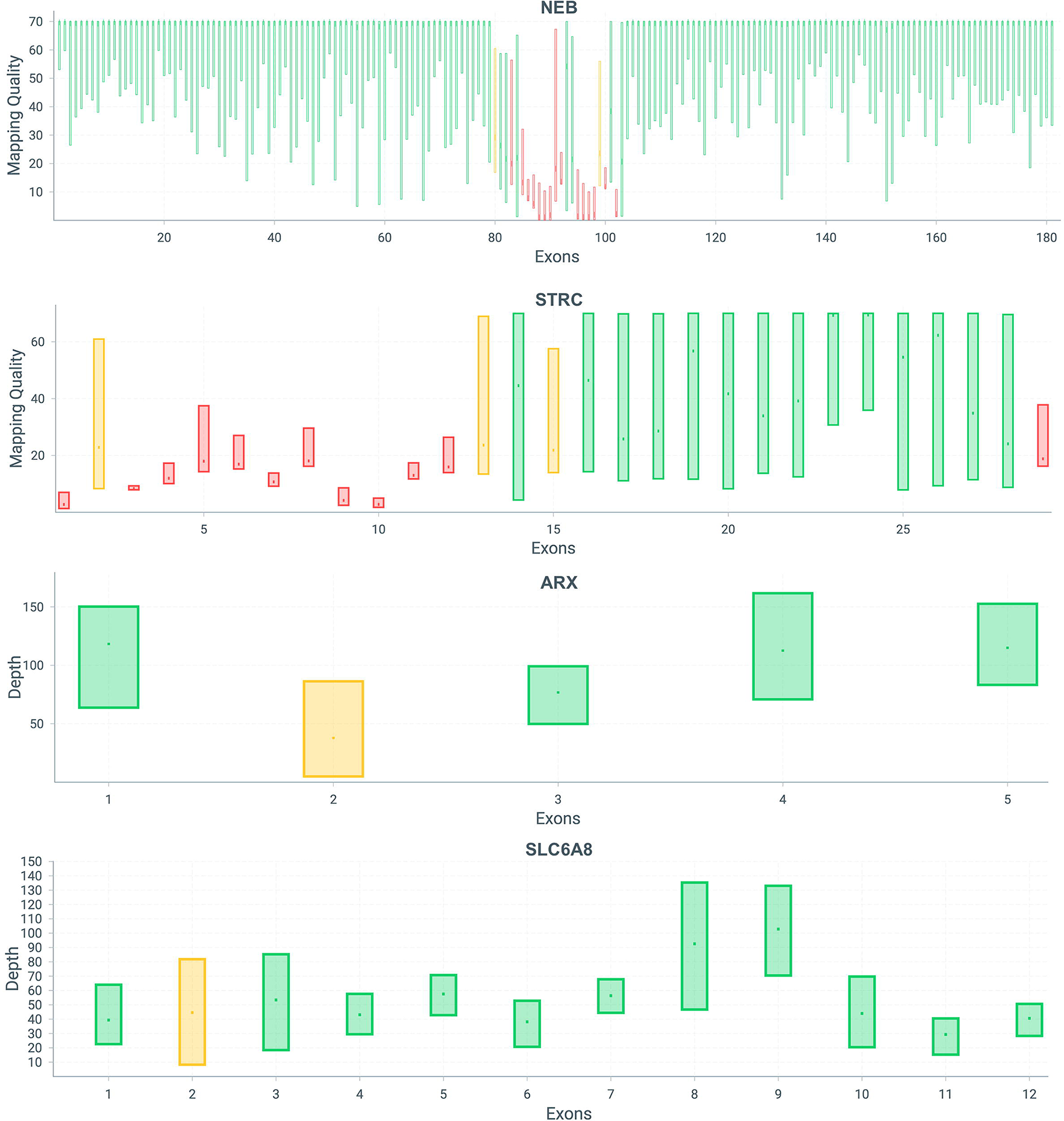
Exome Slicer visualization of four genes with known technically challenging regions, from top to bottom: tandem triplication in Nebulin, *NEB*; a pseudogene in Stereocilin, *STRC;* Poly-Alanine repeat within exon 2 of Aristaless-Related Homeobox, ARX; and GC-rich exons in Creatine Transporter, *SLC6A8.* MQ plots are shown for *STRC* and *NEB*, while coverage plots displayed for *ARX* and *SLC6A8*. Green bars indicate exons with both average MQ ≥ 20 and min DP ≥ 15. Orange bar represents either average MQ < 20 or min DP < 15. Red bars indicate poor exons where both average MQ < 20 and min DP < 15. Each bar in MQ and coverage plots showx minimum and maximum range for each exon (top and bottom of the bar), average is shown by a tick mark in the middle of each bar.

We further assessed the ability of ExomeSlicer to identify technically challenging regions that were characterized using different approaches. First, we targeted 338 known disease genes (**Supplementary Table 1**) using a custom capture (**Supplementary Figure 1**) in three samples, and then performed sequencing to an average overall coverage of ~400x for those samples. We then assessed if technically challenging regions defined as having average MQ < 20 and/or average minimum DP < 15 were comparable using this targeted capture or ExomeSlicer. Indeed, in addition to the homology issues across several exons within the *HYDIN* gene, all 28 poor quality regions in the targeted panel were also identified by ExomeSlicer (**Figure 5** and **Supplementary Table 2**), however, four exons (including *COL5A1* exon 1, see below) exhibited suboptimal quality metrics that would still be identified using more relaxed cutoffs in ExomeSlicer (**Supplementary Table 2**).

**Figure 5.**
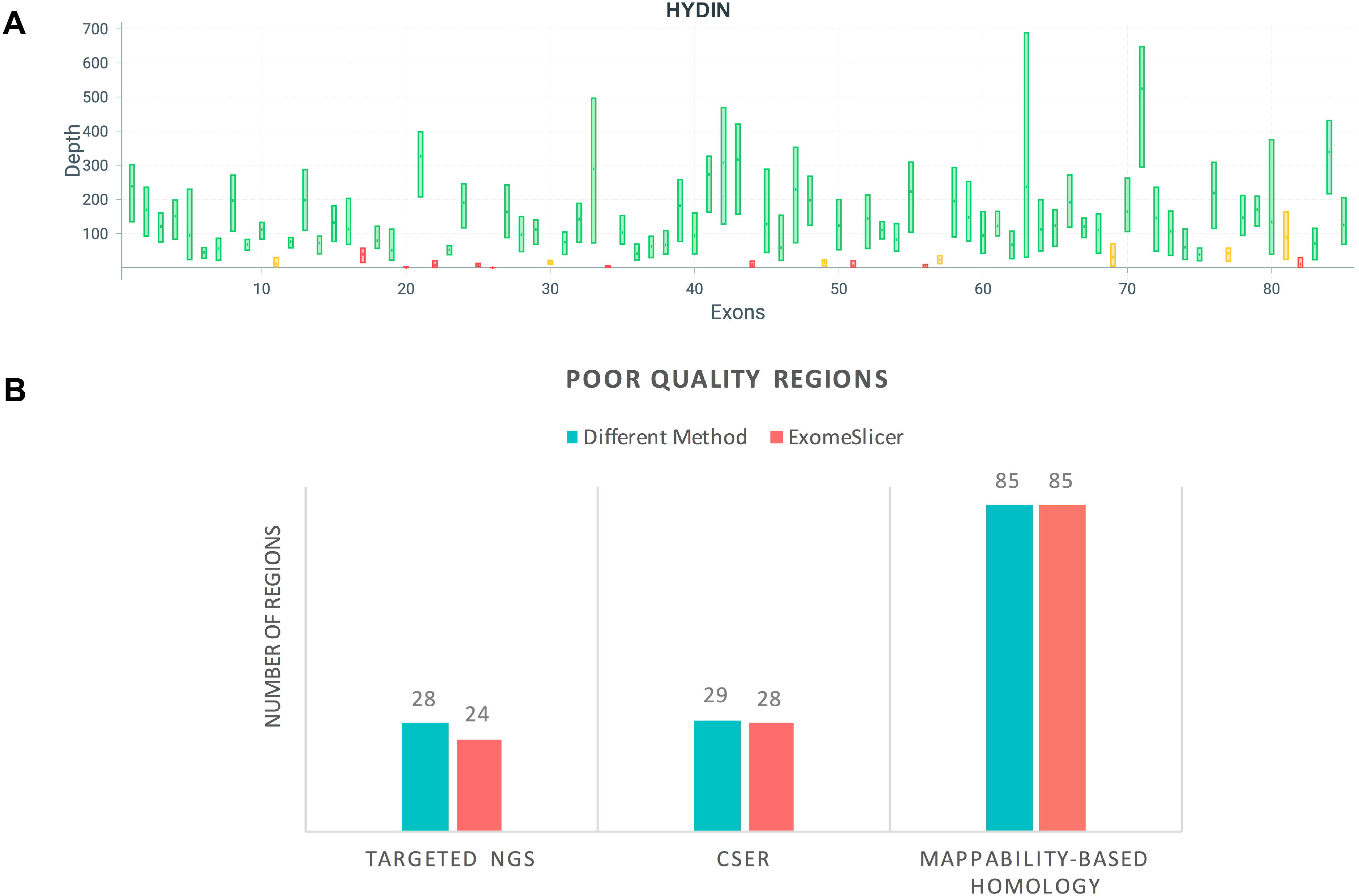
Utility of ExomeSlicer. **A**, ExomeSlicer visualization of *HYDIN*, a gene with significant homology issues. This gene has low quality issues using either a custom-based targeted approach or ExomeSlicer. **B**, Identification of poor quality regions in a targeted NGS panel (with 338 genes) by ExomeSlicer. All 28 regions were called by ExomeSlicer though 4 regions were suboptimal (See **Supplementary Table 2**). Almost all poor quality regions identified by CSER consortium or Mandelker et al were also identified by ExomeSlicer (see **Results** and **Supplementary Tables 3 and 4**).

We also tested if ExomeSlicer would identify the reduced coverage regions recently characterized across 10 centers within the Clinical Sequencing Exploratory Research (CSER) Consortium [16]. With the exception of *COL5A1* exon 1 which again had suboptimal quality, all other regions (28/29) identified in the CSER study were also called by ExomeSlicer (**Figure 5** and Supplementary Table 3). In addition, all homology regions recently detected, in genes with strongest clinical validity, through a different mappability-based approach [25] had low or suboptimal average MQ scores in ExomeSlicer, though these scores had a wide range below normal most likely reflecting different levels of homology within those regions (**Figure 5** and **Supplementary Table 4**).

In summary, the above data demonstrate the utility of ExomeSlicer in predicting regions that cannot be reliably sequenced, most likely due to inherent low complexity sequences (see below).

### Characterization of Exome-wide Poor Quality Regions

Given the adequate coverage achieved using our exome sequencing approach (**Supplementary Figure 2**) across the 1932 samples used to develop ExomeSlicer, we hypothesized that “well-baited” poor quality regions identified by this tool using strict quality cutoffs are most likely to have sequencing complexities obviating appropriate capture, amplification and/or sequencing. However, it is also possible that some low coverage exons are inappropriately baited leading to “false” poor quality annotations.

To distill exome-wide “true” poor quality regions, we first extracted those with either an average MQ < 20, minimum DP < 10, or both (i.e. average MQ < 20 and minimum DP < 10). Then, we only retained those regions that also had at least one capture bait; around 2% (4,515/202,567) of all unique baited exons remained. Despite the presence of one or more baits, their positioning might affect capturing efficiency leading to inadequate coverage. Therefore, we calculated the number of bases covered by bait(s) for each exon, and binned the poor quality regions based on percentage of covered bases (**Figure 6A**). Interestingly, compared to “good quality” regions, which do not meet the above MQ and DP cutoffs (n=198,052), a significant proportion of the 4,515 regions had baits covering less than 70% of exon bases (47% poor quality versus 8% good quality regions, P=0.0016; **Figure 6B**). There was no statistically significant difference between the proportions of poor and good quality regions with over 70% baited bases (P=0.51). Therefore, we only retained the subset of poor regions with more than 70% of their bases covered by bait(s) and deemed as true “poor quality”. They comprised 1.1% (n=2,278) of all baited exons (**Supplementary Table 5**).

**Figure 6.**
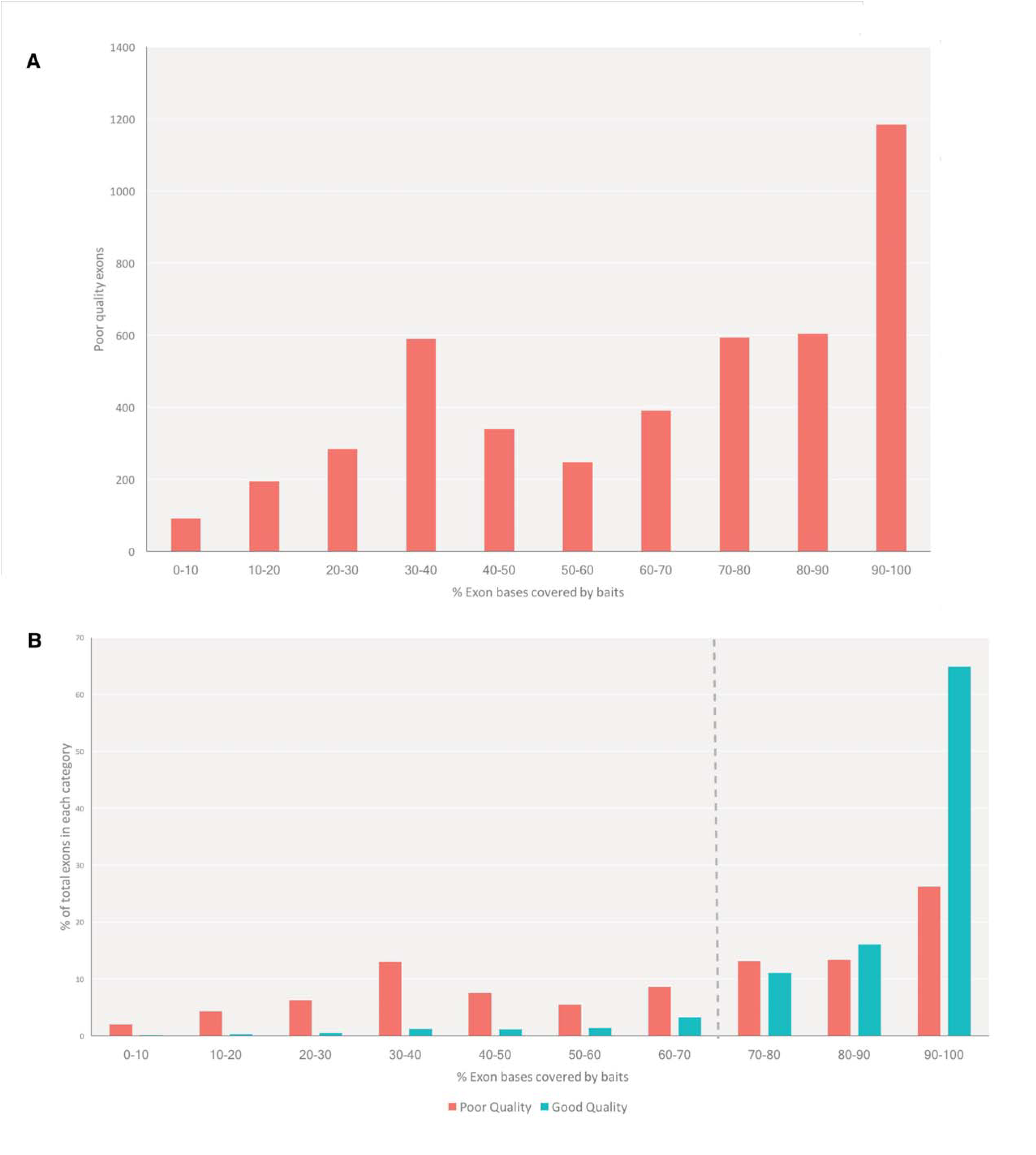
Analysis of baits positioning in targeted regions within the exome. Only regions with at least one bait were included in this analysis. **A**, Distribution of poor quality regions (see **Results** section) based on the percentage of bases covered by baits (X-axis). **B**, Proportions of poor or good quality regions stratified by the percentage of based covered by baits.

We further characterized those regions. We determined their GC content and compared to that of good quality regions of similarly baited (70%) bases (n=175,274). As expected, the former regions had significantly higher GC content (median GC% is 0.62 for poor versus 0.49 for good quality regions, P < 2.2e-16; **Figure 7A**). Of the 2,278 poor quality regions, 846 had mapping quality issues (MQ < 50), and as such had significantly higher BLAT hits (P < 2.2e-16) compared to those with MQ > 50 (**Figure 7B**). The remaining 1,431 regions mostly suffered from low coverage.

**Figure 7.**
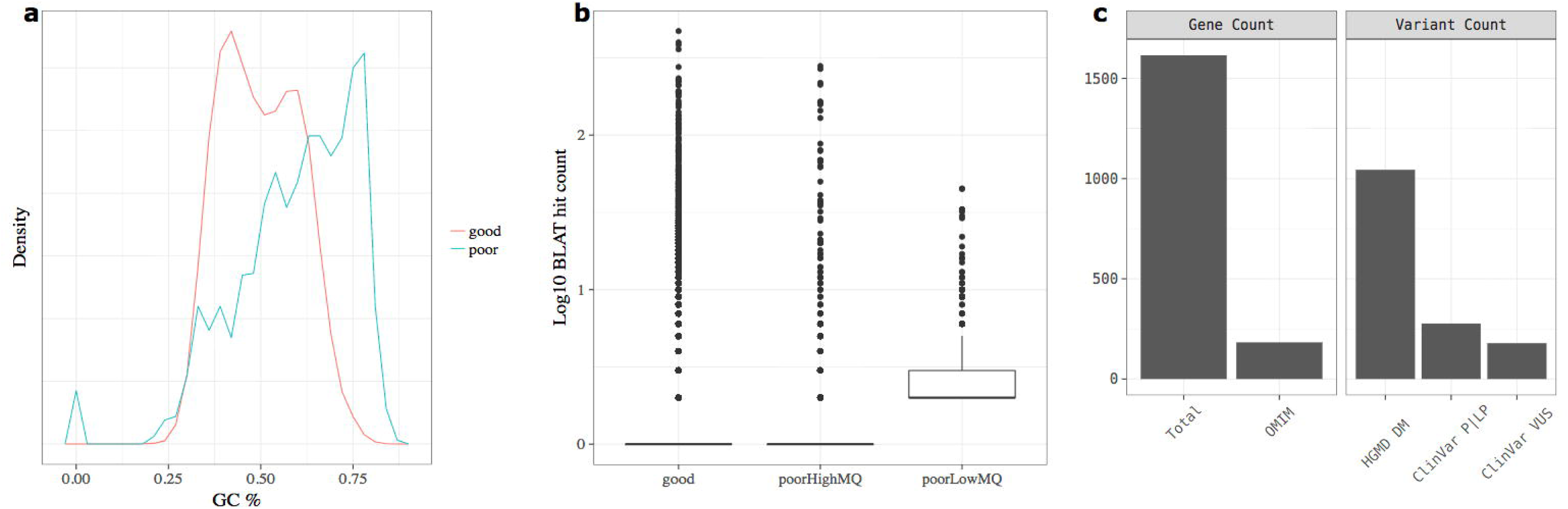
Characterization of exome-wide poor quality regions. This analysis included regions with > 70% of exon bases covered by baits and the MQ and DP cutoffs explained in **Results** section. **A**, Distribution of poor and good quality regions based on percentage GC content (Median GC% is 0.62 and 0.49 in poor and good quality regions, respectively, P < 2.2e-16). **B**, Log10 BLAT hit count of good quality and poor quality regions. The latter was further divided into poorHighMQ (MQ >50) and poorLowMQ (MQ < 50). All poorLowMQ regions had at least 2hits (Median BLAT hits was 2 for poorLowMQ versus 1 for either good or poorHighMQ, P < 2.2e-16). **C**, The 2,278 poor quality regions affected 1,615 genes including 183 OMIM genes. They also contained 1,044 disease mutations (DMs) in the Human Gene Mutation Database (HGMD). In addition, 277 pathogenic/likely pathogenic (P/LP) variants and 179 variants of uncertain clinical significance (VUS) in ClinVar were identified in those regions.

Finally, the poor quality 2,278 regions affected 1,615 genes, 183 of which have been linked to human disease according to the Online Mendelian Inheritance in Man (OMIM) database. Importantly, those regions contained 1,044 disease mutations (DMs) in the Human Gene Mutation Database (HGMD), in addition to 277 pathogenic or likely pathogenic variants and 179 variants of uncertain clinical significance (VUS) in ClinVar database (**Figure 7C**).

## Discussion

Exome-based panels have become an attractive strategy for multigene panel testing in clinical laboratories. This approach reduces the interpretation challenges (relative to WES) while enabling dynamic gene content updates and cost-effective reflex to WES if needed (both are limitation of targeted gene panels). However, there are currently no standards, guidelines, or resources – like our newly developed ExomeSlicer tool – to support the clinical development and implementation of exome-based panels. We have therefore highlighted important considerations associated with using this approach including gene content, exome capture, coverage issues, Sanger fill in, analytical validation and bioinformatics filtration. We also demonstrate the utility of ExomeSlicer and characterize exome-wide technically challenging regions affecting over 1600 human genes.

Although we outline a gene selection strategy, we did not discuss the minimum number of genes at which the exome-based approach should be considered. Sequencing of 5-10 genes might be more cost-effective using a targeted than an exome-based approach. However, this also has to be balanced with the laboratory’s sample volume and existing infrastructure. It might be more disruptive and, therefore, more costly to use a targeted capture and a small-scale sequencing platform if the lab’s main testing volume involves an exome capture and correspondingly high sequencing throughput. Therefore, each laboratory should make a decision about the best platform to use based on cost and their specific needs.

Needless to emphasize, appropriate depth of coverage of the relevant regions of interest (ROI) should be carefully assessed before an exome-based approach is used. Given the large number of targets in an exome, the overall coverage is likely to be lower at a given sequencing throughput. This might in turn lead to more regions falling below the necessary coverage for reliable variant detection, thus leading to an increased burden of Sanger sequencing. Alternatively, deeper sequencing can be achieved though the cost will be higher. Fortunately, current exome kits, including the Agilent CRE kit (**Supplementary Figure 2**), are optimized to enrich for coding regions in general and specifically in disease genes. However, there are two important considerations. First, low complexity, such as pseudogenes, repeats, and GC-rich, regions will often have no or inadequate good quality coverage regardless of capture density or sequencing throughput. Second, because a large number (> 100) of genes is usually involved in exome-panels, more of the above technically challenging regions, which labs are often not familiar with, are expected. Consequently, this will significantly increase the Sanger sequencing fill in burden associated with this approach (**Figure 3**).

Using empirical NGS exon level quality metrics, namely MQ and DP, from 1932 clinical WES samples, we show that ExomeSlicer can support the development of exome-based panels through upfront identification of technically challenging regions, thus guiding ancillary assay development, and reducing post hoc additional analysis, including Sanger sequencing, burden. Because of their inherent low sequence complexities, we show that such regions are not specific to the exome-based approach and should thus be carefully assessed even when a custom-based targeted approach is being used (**Figure 4** and **5**). We do not rule out the possibility that ExomeSlicer might identify false positive low coverage regions due low number or absence of baits or their inadequate positioning. However, this tool displays this information (i.e. number of baits per exon and percentage of exon covered by baits) for the user to distinguish between this and a true poor quality region. In addition, users are encouraged to perform more analysis (such as BLAT, literature search, visual inspection) on any identified regions before considering an ancillary fill in strategy. More important, ExomeSlicer is unlikely to have a high false negative rate, if any. Regions falling below adequate DP and MQ across a large number of samples are more likely to be (or at least equally) identified by ExomeSlicer than by a custom-based targeted approach.

We have introduced stringent metrics (average MQ < 20 and/or average minimum DP < 10x and > 70% of bases covered by baits) to distill exome-wide 2,278 regions with inherent low sequence complexity issues rendering them technically inaccessible to NGS in general. Those regions had significantly higher GC content and, for a subset (n=846), retrieved more BLAT hits indicative of high homology issues (**Figures 6** and **7**). Interestingly, those regions comprised 1.1% of all baited exons in the exome, and are likely to be an underestimate since, we believe, our quality cutoffs were too stringent. Additionally, we cannot rule out that other regions, with less 70% of their exon bases covered by baits, might also have sequencing issues. Intriguingly, those regions were contained in a large number of genes (n=1,615) including 183 that have been associated with human disease, and harbored over 1000 disease mutations in HMGD and over 450 variants of potential clinical significance in ClinVar (**Figure 7**). It is possible that some of those variants are technical false positives calls due to inaccurate sequencing quality. Alternatively, they can be true pathogenic variants identified through different approaches, but can be missed by exome sequencing and NGS in general. Either scenario supports the strong possibility of false positive and/or false negative variant calls by NGS/WES in those regions, thus leading misdiagnoses in a potentially large number of patients. Finally, 1,432 impacted genes are not linked to disease yet and should, therefore, be carefully approached by researchers to avoid inaccurate annotations and/or functional analysis.

In summary, our work provides a comprehensive overview and a novel resource, ExomeSlicer, for the validation and clinical implementation of exome-base panels. In addition, our characterization of the exome-wide poor regions represents a valuable resource for clinicians and researchers using whole exome sequencing.

## Figure Legends

**Supplementary Figure 1**. Gene selection strategy. Epilepsy is used as a disease example. In the first step, genes that are non-syndromic or wherein epilepsy in the main diagnostic feature are assessed for validity using the ClinGen framework. Only genes with strong clinical validity are included. A list of 100 genes, included on an epilepsy exome-based clinical panel at The Children's Hospital of Philadelphia, is shown.

**Supplementary Figure 2**. Coverage comparison between exome-based (Agilent CRE kit, n=1932) and custom-based targeted (n=400) enrichment of 338 disease genes. Those genes were sequenced to 100x (exome) or 400x (targeted) depth of coverage. ***Top***, comparison of coverage across all 338 genes and a selection of disease panels. ***Bottom***, comparison of coverage across representative 22 epilepsy disease genes.

